# The dynamic ontogenetic patterns of adaptive divergence and sexual dimorphism in Arctic charr

**DOI:** 10.1101/2021.01.15.426104

**Authors:** Marina De la Cámara, Lieke Ponsioen, Quentin J.B. Horta-Lacueva, Kalina H Kapralova

**Author notes:** Corresponding author: Marina de la Cámara, Correspondence.

## Abstract

Arctic charr (*Salvelinus alpinus*) in lake Thingvallavatn (Iceland) is one of the most iconic examples of post-glacial adaptive divergence, resulting in four ecomorphs that diverge along the ecological benthic-limnetic axis (bottom lake *versus* open water feeders), and are distinct both phenotypically and genotypically. Here, we used geometric morphometrics tools on a common garden setup to determine the factors responsible for genetically based shape variation during the post-embryonic ontogeny of two morphs that represent the benthic-limnetic axis: the small benthic (SB) and the planktivorous (PL). This experiment uses pure crosses and F1 reciprocal hybrids between the two morphs, and includes the onset of sexual maturation, offering an excellent opportunity to explore the genetic component of adaptive divergence and the role of sexual dimorphism in this scenario. We found that growth is the main driver of shape variation across time and provided evidence of a genetically-controlled ontogenetic shift that gives rise to the limnetic morph. Additionally, our results indicate that the onset of sexual maturation triggers differences both in sex ontogenetic trajectories and in static shape variation at different time points, likely dissipating the canalisation for traits traditionally associated with benthic-limnetic adaptations.

## INTRODUCTION

Polymorphic and sexually dimorphic populations can be the outcome of natural, sexual and/or ecological sexual selection acting on available phenotypic variation. Recent adaptive radiations have served the literature to understand the origins of polymorphic populations (Schluter, 2000; Losos & Schneider, 2009). However, the interplay between adaptive and sexual traits and their dynamics during ontogeny has been overlooked in the study of evolutionary mechanisms leading to adaptation and speciation (but see Bolnick & Doebeli, 2003; Butler et al. 2007; Cooper et al., 2011; Parsons et al., 2015). Here, we determine the factors responsible for genetically-based shape variation during the post-embryonic ontogeny of a textbook example of polymorphic population: the Arctic charr ecomorphs in lake Thingvallavatn (Iceland), in a long running common garden setup which includes the onset of sexual maturation.

Archetypal examples of adaptive radiations are Darwin’s finches, African cichlids, Hawaiian spiders, *Anolis* lizards, three-spined sticklebacks and salmonids (Grant & Grant, 2002; Meyer 1993; Gillespie, 2004; Losos & Schneider, 2009; Jones et al., 2012; Snorrason & Skúlason, 2004, respectively). These radiations have a common denominator: an ancestral population presents enough genotypic variation to diversify into two or more subpopulations which inhabit a variety of environments, in response to certain selective pressures (i.e. changes in density, resource availability, predators, pathogens or abiotic factors that affect fitness). Distinct traits are then selected to facilitate the exploitation of those environments.

Arctic charr in Thingvallavatn represents a special case of adaptive radiation called resource polymorphism, where distinct phenotypes evolve based on different resource utilisation. Within this lake, four morphs have diverged -both phenotypically and genotypically-, segregating along the water column: two bottom-feeders (i.e. a small and a large benthic) and two limnetic-feeders (i.e. a planktivorous and a piscivorous morph) (Jonsson et al., 1988; Snorrason et al., 1989; Malmquist et al., 1992; Sandlund et al., 1992; Kapralova et al., 2011; Kapralova et al., 2013; Guðbrandsson et al., 2019). This benthic-limnetic adaptive diversification is a widespread evolutionary strategy in teleosts, showing similar phenotypic adaptations (i.e. benthic morphs generally present subterminal jaws, blunt snouts, deep bodies, smaller eyes and less gill rackers, whereas limnetic morphs have terminal jaws, pointed snouts, slender bodies, bigger eyes and more gill rackers) (Sandlund et al., 1992; Snorrason et al., 1994; Berra 2001; Blake et al., 2005; Husley et al., 2013).

Sexual dimorphism in a context of adaptive radiation might arise prior or post ecological diversification, or both (Van Dooren et al., 2004; Aguirre et al., 2008). Ancestral sexual dimorphism of a clade can either facilitate or constrain subsequent adaptive divergence (e.g. Aguirre et al., 2008; De Lisle & Rowe, 2015). Alternatively, theoretical studies suggest that sexual traits evolving after sympatric radiations should be exclusive within each ecomorph, unless assortative mate choice occurs before trait divergence starts (Slaktin 1983; Van Dooren et al., 2004; Parsons et al., 2015). Regardless on its origin, sexual dimorphism has shown major effects in the morphological variation of several adaptive diversification systems, such as *Anolis* lizards, *Notophtalmus* salamanders, dwarf chameleons, endemic roundfins and African cichlids (Butler & Losos, 2002; De Lisle & Rowe, 2017; Da Silva & Tolley, 2012, Pfaender et al., 2011; Parsons et al., 2015).

Besides sticklebacks (e.g.: Reichmen & Nosil, 2004; Kitano et al., 2007; Aguirre et al., 2008; Brener et al., 2010; Cooper et al., 2011; McGee & Wainwright, 2013), our knowledge on the interplay between sexual dimorphism and ecological adaptation in northern freshwater fish radiations is still limited. Nevertheless, a few early studies on salmonids pointed towards complex relationships between adaptive and secondary sexual traits. For instance, a stunted population of Arctic charr in Norway showed that sexual dimorphism overrode morphological changes associated with benthic-limnetic ontogenetic shifts (Bjøru & Sandlund, 1995). More recently, a common garden experiment on Arctic charr from three different Finnish lakes showed morphological differences between populations outside the breeding season in contrast to major effect of sexual traits during breeding season (Janhunen et al., 2008). These studies suggest that sexual dimorphism is not only complex but also variable during the lifecycle in allopatric populations. Previous studies on Arctic charr have found morphological differences related to sympatric benthic-limnetic diversification in early stages of their ontogenies, mainly associated to growth, indicating that these traits may be canalised across time (Skúlason et al., 1989; Kapralova et al., 2015), although the degree of canalisation appears to variable in different lakes (Parsons et al., 2011). Yet, the genetic component of how and when these adaptive traits evolve in later ontogenetic stages of Arctic charr, and the role of sexual dimorphism in this scenario is still unknown.

Common garden experiments present a unique opportunity to standardize environmental cues and elucidate whether certain phenotypic differences (that may be associated with adaptive traits) have a purely genetic component (reviewed by De Villemereuil et al., 2016). For this study, offspring of wild small benthic (SB) and planktivorous (PL) morphs from Thingvallavatn and their reciprocal hybrids were reared in common garden up to 36 months after hatching and phenotyped at four time points during ontogeny (i.e. 12, 18, 24 and 36 months after hatching). Their comparable sexual maturation times (i.e. from 2 (males) to 4 (females) years in SB and 3.5 years in PL) and overlapping spawning periods (Jonsson et al., 1988), facilitated this long-running crossing design.

In a common garden setup, the generation of hybrids helps us gain insight in the evolutionary mechanisms partaking in the diversification continuum (e.g. Skúlason et al., 1989; Skúlason 1993; Grant & Grant, 1994; McGirr & Martin, 2019). In a process of ecological diversification, hybridisation between ecomorphs may be understood as speciation reversal. However, it can also provide an external source of variation that may be adaptive if selective pressures change, leading to adaptive introgression and hybrid speciation (Abbott et al., 2013, and see e.g., Pardo-Díaz et al., 2012; Meier et al., 2017). Although recombination between closely related species results in genotypic mosaicism (in haplodiploid hybridisation), potentially successful phenotypic outcomes can result in intermediate, transgressive and parental-like phenotypes (e.g. Seehausen, 2004; Elgvin et al., 2017; Skúlason et al., 1989, respectively). In sum, exploring the phenotypic outcome of hybrids sheds light on the evolutionary potential of the population.

In Thingvallavatn, gene flow between SB and PL Arctic charr morphs is putatively negligible (Jonsson et al., 1988; Skúlason et al., 1989; Kapralova et al., 2011), despite their recent evolutionary divergence and similarities in life history traits. Nevertheless, a recent common garden experiment on the same morphs from Thingvallavatn showed different patterns of trait covariance between pure crosses and F1 hybrids (rather than mean values of those traits) which may result in lower fitness for the hybrids (Horta-Lacueva et al., *in press*). Regardless of the strength of pre- or post-zygotic reproductive barriers, it is possible to generate both SBxPL and PLxSB crosses in controlled conditions and study their phenotypic outcome throughout their lifetime.

A recently evolved population with a relatively clear evolutionary history, a long-running common garden experiment with pure crosses and hybrids, including the onset of sexual maturation offers a unique framework to explore the genetic component of adaptive divergence and the role of sexual dimorphism in their phenotypes. For this study, we aimed to investigate the factors driving genetically-based shape variation throughout ontogeny of two Arctic charr ecomorphs and their hybrids. More specifically, we ask the following questions: are traits associated with benthic-limnetic adaptations present in a common garden setup? If so, is there evidence for trait canalisation? Are there signatures of sexual dimorphism? Do adaptive and secondary sexual traits overlap, or alternatively, do they affect different dimensions of shape variation? Despite its origin, is sexual dimorphism constraining or facilitating adaptive speciation? How do these traits differ in the reciprocal hybrids? Are there differences in ontogenetic trajectories between cross types and sexes? When do sex and adaptive traits emerge during ontogeny and what is their relative weight at each separate time point?

## MATERIAL AND METHODS

### Data collection

#### Generation of crosses and rearing

Wild adult specimens of the planktivorous (PL) and small benthic (SB) morphs were collected by laying gillnets overnight at a spawning site shared by both (Svínanesvík, 64°11’24.6”N; 21°05’40.5”W) in the beginning of October 2015. 31 mature specimens were crossed either within the same morph (i.e. pure crosses, PLxPL and SBxSB) or reciprocally with the alternative morph (i.e. reciprocal hybrid crosses, PLxSB and SBxPL), generating 19 full-sibling families (as five individuals were involved in two crosses) (see crossing design in Supplementary material (SM), table S1). Fertilised eggs from each family were placed in separate mesh cages in an EWOS hatching tray (EWOS, Norway) at 4.1±0.2°C in the aquaculture facilities of Hólar University College, Sauðárkrókur, Iceland. After hatching and first feeding, families were transferred into separate buckets (35cm deep, ⌀ 29cm) on the same running water flow and commercial dry pellets were administrated manually on the surface in all buckets to ensure common feeding conditions. Fish were phenotyped at four time points throughout ontogeny: at 12, 18, 24 and 36 months after hatching. At each time point, specimens were anaesthetized with 2-phenoxyethanol (Pounder et al., 2018) with a variable dosage according to the fish individual reactions, prior to phenotyping. At 12 months specimens were photographed. At 18 months all fish were PIT- tagged, weighted, photographed and placed together in two aquaculture tanks with similar densities. At month 24, fish were again weighted, photographed, their sex was determined if they had reached sexual maturation, and at month 36 they were photographed and sex determined if possible. Since fish were PIT- tagged at month 18, individual information collected at later time points could be traced back only to then, thus no sex data were available for month 12. Mortality throughout the experimental setup was generally low (less than 10%), and higher levels of mortality were not attributed to any specific cross type or family.

#### Photographing

Photos of each individual were taken on their left lateral side with a fixed digital camera (Canon EOS 650D and 100mm macro lens) at the four different time points along with a piece of graph paper or a ruler for scaling. Fish tend to naturally bend as they are positioned on a rigid flat surface while the photos are being taken, which confound biologically meaningful shape changes (Valentin et al., 2008). We corrected for potential bending effects by placing 5 equidistant landmarks along the lateral line of each fish and implementing the unbending tool in tpsUtil (Rohlf 2016).

#### Landmarking

We placed 16 landmarks and 5 semilandmarks on each photo using tpsDig (version 2.26, Rohlf, 2016) (**Fig. 1**), based on previous geometric morphometrics literature on Icelandic Arctic charr (Adams & Huntingford, 2004; Parsons et al., 2010). For scaling, two additional landmarks were placed on graph paper or a ruler. Landmarking of the data was conducted by the same person (LP). Thirty random individuals from different cross types and families were landmarked three times, obtaining a high repeatability (p < 0.05) which ensured the robustness of the data.

**Figure 1.**
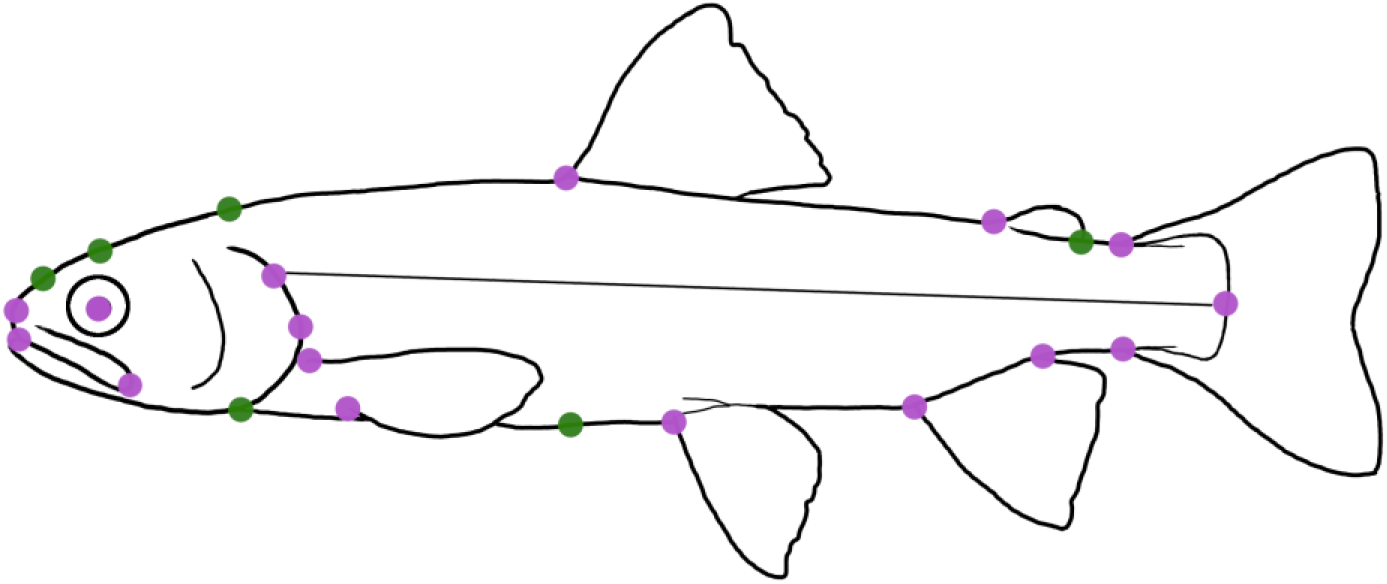
Landmark (violet) and semilandmark (green) configuration used for geometric morphometrics analyses. Landmarks taken for scaling and unbending are not shown.

### Analysis of shape

Raw coordinates in .tps format were imported in R and subsequent analyses were conducted with the geometric morphometrics package *geomorph v3.3.1.* and *RRPP v0.6.1* (Collyer & Adams, 2018; Adams et al., 2020; Collyer & Adams, 2020).

Outlier examination was conducted by looking at Procrustes distance of each landmark configuration to their mean shape, grouped by month and cross type. Configurations falling on the upper quantile of the distribution represented 2.18% of the full data set and were inspected individually, showing either landmark displacements or slightly open jaws. After outlier removal, Partial Generalized Procrustes Superimposition was performed on raw data and the resulting Procrustes coordinates were used in downstream analyses (see **Table 1** for number of individuals used at each time point). During Procrustes superimposition, each landmark configuration is translated, scaled and rotated to minimise shape differences among them. Centroid size, which is the square root of the sum of squared distances of the landmarks to the centroid, is extracted when scaling.

**Table 1.**
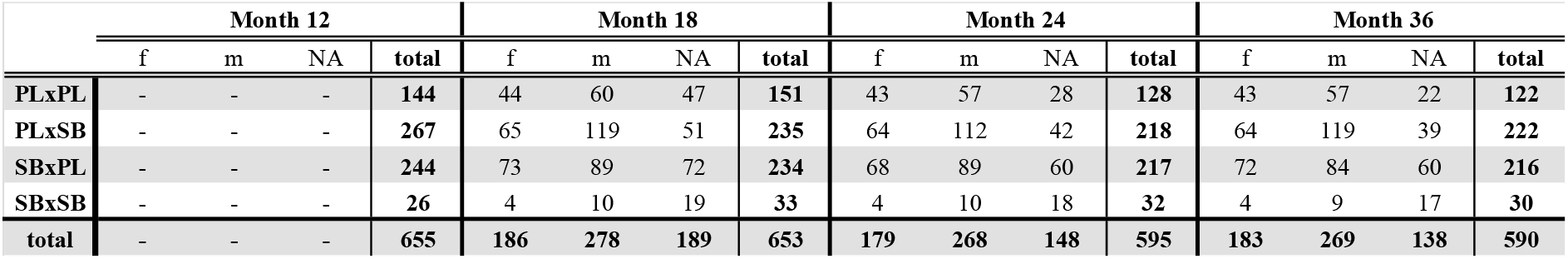
Number of individuals used for geometric morphometrics analyses (outliers removed) per cross type and sex at each time point.

|  | Month 12 |  |  |  | Month 18 |  |  |  | Month 24 |  |  |  | Month 36 |  |  |  |
| --- | --- | --- | --- | --- | --- | --- | --- | --- | --- | --- | --- | --- | --- | --- | --- | --- |
|  | f | m | NA | total | f | m | NA | total | f | m | NA | total | f | m | NA | total |
| PLxPL | - | - | - | 144 | 44 | 60 | 47 | 151 | 43 | 57 | 28 | 128 | 43 | 57 | 22 | 122 |
| PLxSB | - | - | - | 267 | 65 | 119 | 51 | 235 | 64 | 112 | 42 | 218 | 64 | 119 | 39 | 222 |
| SBxPL | - | - | - | 244 | 73 | 89 | 72 | 234 | 68 | 89 | 60 | 217 | 72 | 84 | 60 | 216 |
| SBxSB | - | - | - | 26 | 4 | 10 | 19 | 33 | 4 | 10 | 18 | 32 | 4 | 9 | 17 | 30 |
| total | - | - | - | 655 | 186 | 278 | 189 | 653 | 179 | 268 | 148 | 595 | 183 | 269 | 138 | 590 |

Because body weight was missing at time points 12 and 36 and could not be used as an alternative measure to centroid size, we calculated their correlation. Centroid size may not represent an adequate proxy of body size for multiple reasons, for example an uneven placement of the landmarks along the shape outline (Collyer et al., 2020). This might have occurred in our data set, since Arctic charr has a fusiform body and we placed a considerable proportion of landmarks on the head. Therefore a high correlation between body weight and centroid size is a reliable indicator that centroid size may be a good proxy for body size. Effectively, we found a correlation of 98.6% (Pearson’s correlation coefficient, p < 0.05) on the log-log regression of both variables, and therefore *log(Csize)* was used as a proxy for body size in subsequent analyses. One-way ANOVA followed by post-hoc Tukey tests were performed in base R to study *log(Csize)* between cross types and sexes at each time point.

Statistical analyses on shape were performed at two different levels: on specimens which reached sexual maturation during the experimental set up (1) and thus individual sex information could be traced back to month 18 and studied across time; and on a full data set of specimens (2) to study the dynamic patterns of shape variation across four time points (i.e. month 12, 18, 24 and 36) with no sex information and hence heterogeneity in the sexual maturation state of the specimens. For both datasets, the static patterns of shape variation were additionally explored at each time point separately.

For the sexed specimens (1), we performed mixed-model MANOVAs with *geomorph::procD.lm* to examine the effects of *log(Csize), month, sex* and *cross type* with nested families (*cross/family*) (including all possible interactions) on *shape* (i.e. and here after, Procrustes coordinates). We used the *RRPP::pairwise* function to assess differences in shape between group means, variances (i.e. morphological disparity) and vector correlations. The dynamics of morphological disparity can be highly informative when studying the ontogenetic changes in morphs involved in adaptive divergence, since a reduction of morphological disparity within each group across time can be a proxy for canalisation of certain traits (e.g. Lazić et al., 2014; Haber & Dworkin, 2016; Vučić et al., 2019).

Phenotypic Trajectory Analysis (PTA) was implemented in *geomorph* to study ontogenetic trajectories of sexual dimorphism where each time point (month 18, 24 and 36) was used as a trajectory point. Due to unbalanced number of sexed individuals among cross types and families (see **Table 1**, and SM1 Table 1), we decided to pool the four cross types together and study intraspecific sex trajectories as a whole. We explored pairwise differences in location, length (i.e. amount of shape change), directionality and shape between both trajectories. For the same data set, MANOVAs and pairwise tests were performed separately on the different time points. We used *geomorph::plotAllometry* for visual interpretation of static allometry patterns and Homogeneity of Slopes (HOS) tests were conducted by comparing models of common (*shape ~ log(Csize) + sex*) versus unique allometry (*shape ~ log(Csize) * sex*) statistically in order to decide whether to correct for static allometric effects within each time point.

For the full data set (2) we used the same functions as in (1), this time including an additional time point (i.e. month 12) and excluding the *sex* covariate. Increasing sample size and lengthening ontogenetic trajectories allowed for a more complete examination of shape changes associated with growth and ontogenies of pure crosses and F1 hybrids. Again, mixed-model MANOVAs, pairwise tests and HOS test were conducted on separate life stages for this data set.

#### Visualisation

Principal Component Analyses (PCA) on Procrustes coordinates were conducted for visualisation of shape variation at each level. Additionally, we used wireframes to explore shape changes in landmark configurations at the extremes of the most relevant eigenvectors. Ontogenetic trajectories were plotted onto the first two principal components based on the covariance matrix of group means. Trajectories connect mean shape estimates at the different time points for each sex or cross type.

## RESULTS

### Size, month, sex and cross type drive shape variation during ontogeny

We first explored the drivers of shape and their relative weight on the sexed individuals, finding that size, time point, sex, cross type and multiple interaction effects explained shape variation in this experimental setup, as shown in **Table 2**. The relative load of each covariate was analysed by looking at standardised effect sizes (i.e. Z scores). Effect sizes showed that centroid size, month and their interaction had an important effect, suggesting that growth is strongly driving shape changes across ontogeny. The interaction of centroid size with each term except *cross* alone additionally indicated that size plays a key role in shape variation for different sexes and families. On the other hand, *sex* alone had a strong effect on shape, pointing towards a major influence of the onset of sexual maturation, regardless of time point or cross type.

**Table 2.** Significant terms affecting shape of sexed individuals phenotyped at 18, 24 and 36 months after hatching in a mixed-model MANOVA using residual randomization. Error terms updated to account for nested families within cross types.

|  | d.f. | SS | MS | Rsq | F | Z | Pr (>F) |
| --- | --- | --- | --- | --- | --- | --- | --- |
| *log(Csize)* | 1 | 0.078 | 0.0776 | 0.069 | 229.595 | 10.562 | 0.001 |
| *month* | 2 | 0.085 | 0.0427 | 0.076 | 126.196 | 12.387 | 0.001 |
| *sex* | 1 | 0.049 | 0.0489 | 0.044 | 144.58 | 9.644 | 0.001 |
| *cross* | 3 | 0.055 | 0.0183 | 0.049 | 4.061 | 4.484 | 0.001 |
| *cross:family* | 14 | 0.063 | 0.0045 | 0.056 | 13.352 | 17.552 | 0.001 |
| *log(Csize):month* | 2 | 0.009 | 0.0046 | 0.008 | 13.609 | 7.748 | 0.001 |
| *log(Csize):sex* | 1 | 0.025 | 0.0246 | 0.022 | 72.617 | 9.004 | 0.001 |
| *log(Csize):month:sex* | 2 | 0.010 | 0.0049 | 0.009 | 14.612 | 8.224 | 0.001 |
| *log(Csize):month:sex:cross:family* | 99 | 0.103 | 0.0010 | 0.092 | 3.086 | 18.015 | 0.001 |
| Residuals | 1231 | 0.416 | 0.0003 | 0.371 |  |  |  |
| Total | 1356 | 1.121 |  |  |  |  |  |

Interestingly, cross type still had a significant impact on shape even though individuals sampled through a great proportion of their lifespan were pooled together in the analyses, and thus its signal was substantially lower compared to the other terms. Nevertheless, nested families showed the largest Z score both alone and together with *log(Csize)*month*sex* term, which suggests that the effect of individual family trajectories across time is comparable to the overall growth effect.

To better understand the relative effects of each covariate across time, we performed MANOVAs at the three later time points. We found significant effects on shape for *log(Csize), cross, cross:family, sex* and *log(Csize):sex:cross:family* at each time point, and *log(Csize):sex* was only significant at month 24 (**Fig. 2**). Despite both cross and sex being significant across ontogeny, the effect of sex abruptly increased from month 18 to 24 and 36, whereas cross, which had larger load than sex at month 18, greatly decreases at 36. We found males were significantly larger than females at months 18 and 24 (t = 3.134, p = 0.0018 and t = 2.29, p = 0.0013 respectively), but no centroid size differences were found at month 36 (t = −1.479, p = 0.14; see SM2, table S2.2).

**Figure 2.**
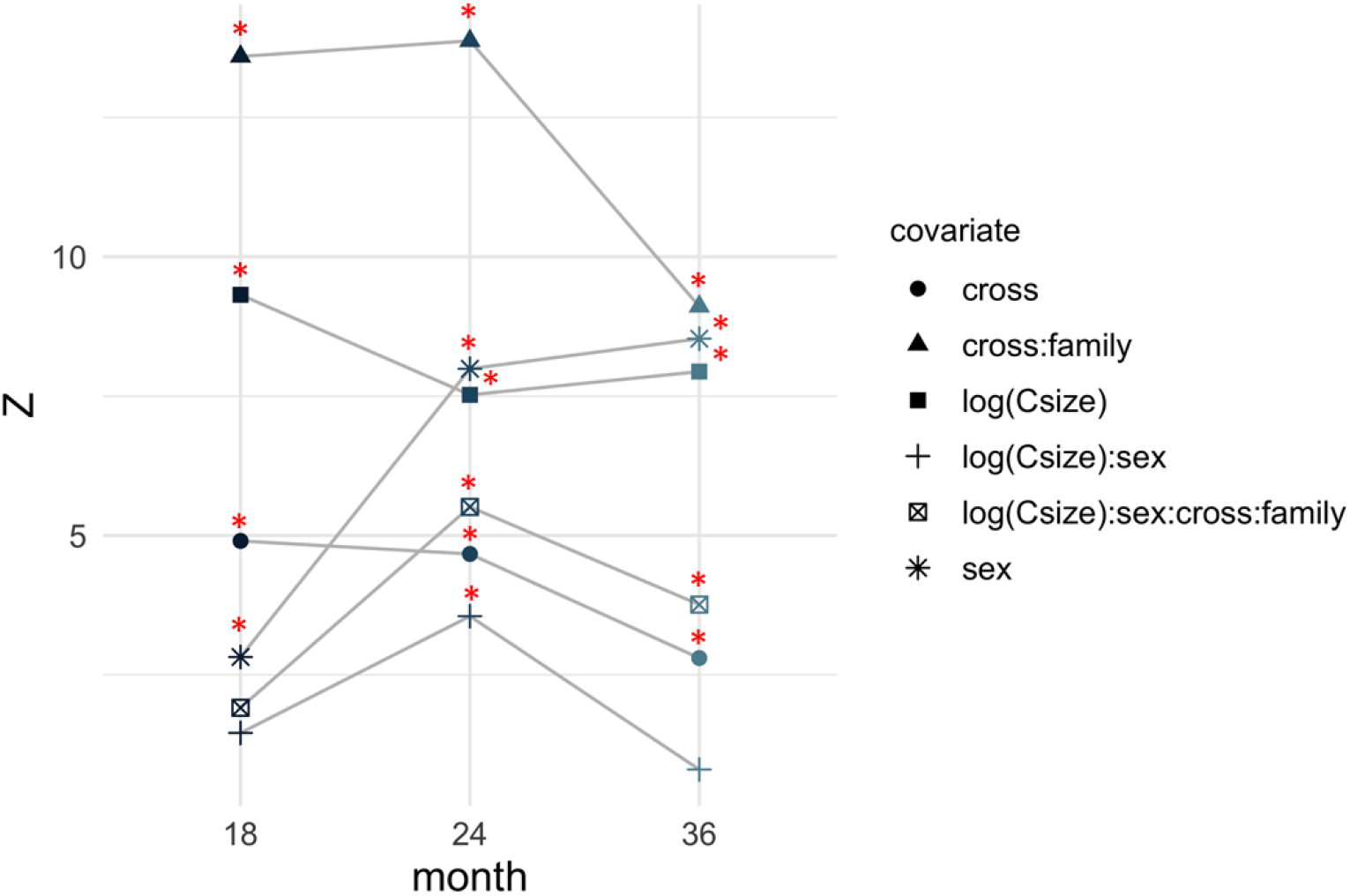
Relative loads of significant covariates in independent MANOVAs at month 18, 24 and 36, represented by Z scores and p-values < 0.05 (red asterisks).

### Juveniles look benthic, adults look limnetic

Considering the relative importance of growth in shape variation across time, we included individuals who were not sexually mature during the experimental set up and hence added a fourth time point at month 12 after hatching. For this model, *log(Csize), month, cross, cross:family* and *log(Csize)* interactions were highly significant, again finding the largest weights in *log(Csize), month, cross:family* and the interaction *log(Csize):month:cross:family* (SM3, table S3).

When fitting explicit reduced models for *log(Csize)*cross/family*, we found significant pairwise differences in means between all months (Pr > d = 0.001), with the largest Z scores for mean comparisons between month 36 and the rest (12-36, Z = 18.31; 18-36, Z = 22.56; 24-36, Z = 19.41). There were significant pairwise differences in morphological disparity between all time points (Pr>d ≤ 0.003) except for 12-18, again with the largest differences between month 36 and the rest (12-36, Z = 23.47; 18-36, Z = 23.35; 24-36, Z = 21.61). As expected from the previous test, cross type had a significant effect. Specifically, when fitting explicit reduced models for *log(Csize)*month*, we found significant pairwise differences in means between each cross (Pr > d = 0.001), and also in variance between SBxPL and PLxPL, and SBxPL and PLxSB (Pr > d = 0.001).

Consistently, Principal Component Analysis (PCA) of Procrustes coordinates (**Fig. 3A**) revealed clustering by month along both PC1 (36.50%) and PC2 (21.57%), although shape variance along the first six dimensions was also examined. Specimens at 12 and 18 months diverged mainly along PC2, although month 18 spanned the distance towards month 24 along PC1. Month 24 had a more scattered distribution in the morphospace, overlapping with both months 18 and 36, and increasing its morphological disparity. Further, a considerable proportion of the morphospace is occupied by specimens phenotyped at month 36, where morphological variability increased markedly relative to previous time points (**Fig. 3B**, and see SM4, table S4).

**Figure 3.**
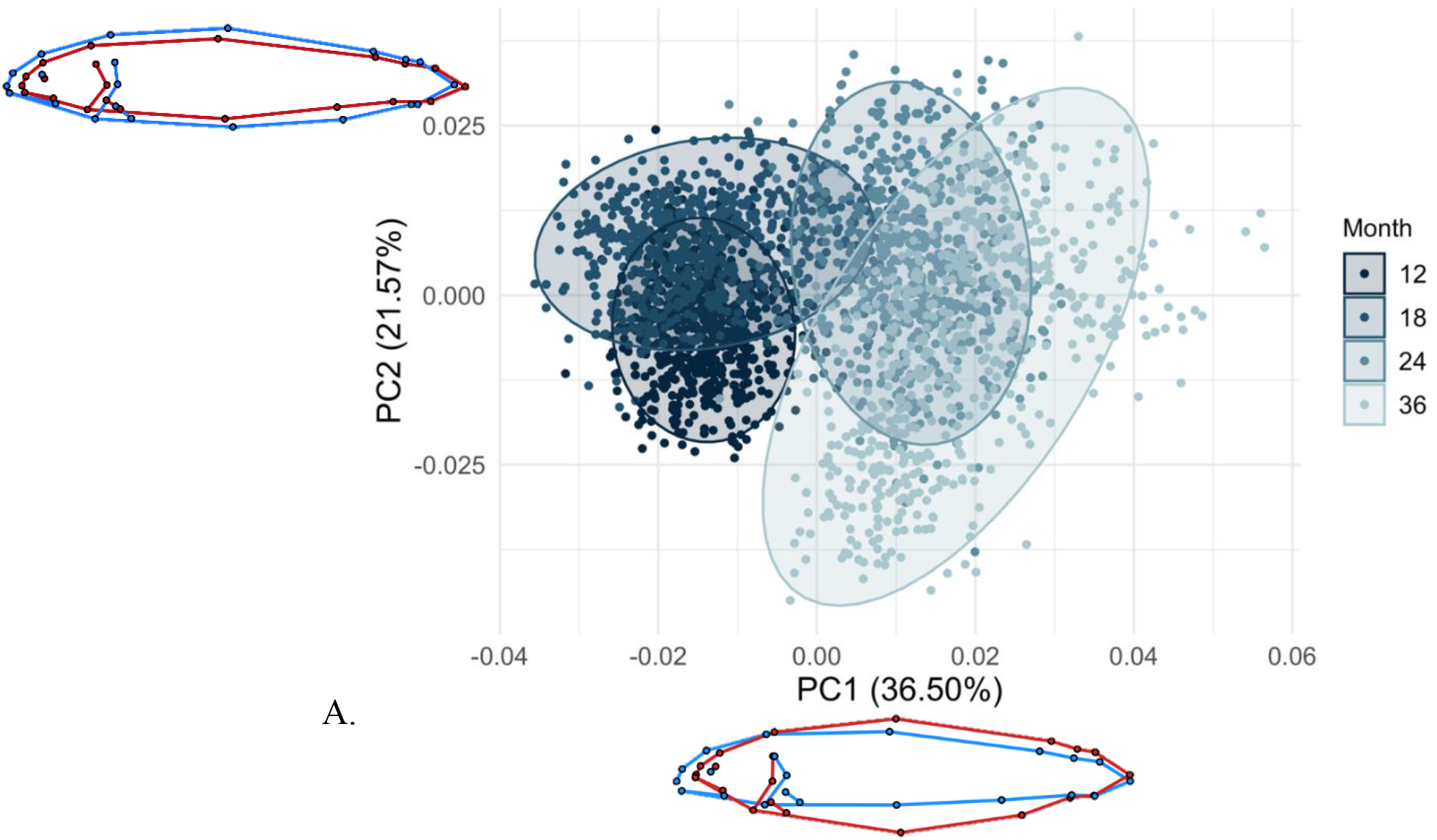
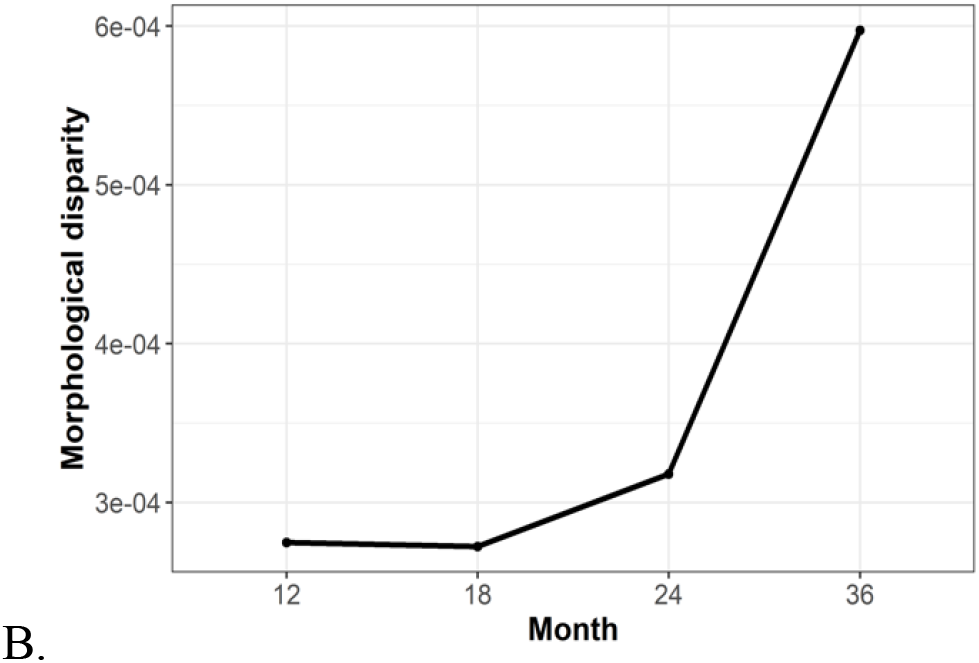
**(A)** Principal Component Analyses of Procrustes coordinates for every specimen. Each point represents one individual. Navy blue represents specimens at month 12, steel blue at 18, medium blue at 24 and light blue at month 36. Shaded, 95% confidence ellipses for each time point. Wireframes depict landmark configurations at maximum (red) and minimum (blue) values for PC1 and PC2. **(B)** Morphological disparity at each time point.

Shape changes along PC1 affect both head and body shape. Younger individuals (i.e. months 12 and 18) have relatively larger heads, rounder snouts, longer upper jaws and slender bodies. On the other hand, individuals at month 36 tend to present relatively smaller heads, pointed snouts and deep bodies, regardless on cross type. Low values of PC2 show much deeper heads and bodies and a shorter caudal peduncle, whereas high values show narrower heads and bodies and longer caudal peduncles.

### Sexual maturation triggers differences in sex ontogenetic trajectories and shape variation at each time point

Since the onset of sexual maturation occurred during the experimental setup and had an overall important impact on shape (**Table 2**), we explored the ontogenetic trajectories of potential sexual dimorphism.

Phenotypic Trajectory Analysis (PTA) (**Fig. 4**) showed significant differences in trajectory location, directionality and shape between sexes (p = 0.001, Pr > angle = 0.001 and Pr > d = 0.001 respectively), although absolute path distances did not differ (Pr > d = 0.94). At month 18, mean shapes of both sexes laid practically at the same point in the morphospace. However, at month 24, they diverged along PC2, and differences increased towards month 36, where the relatively large morphological disparity seemed to be explained by -at least- a very marked sexual dimorphism. Additionally, morphological disparity in males is not only larger, but seems to increase at a higher rate than morphological disparity in females (SM4, Table S4, fig. S4.1).

**Figure 4.**
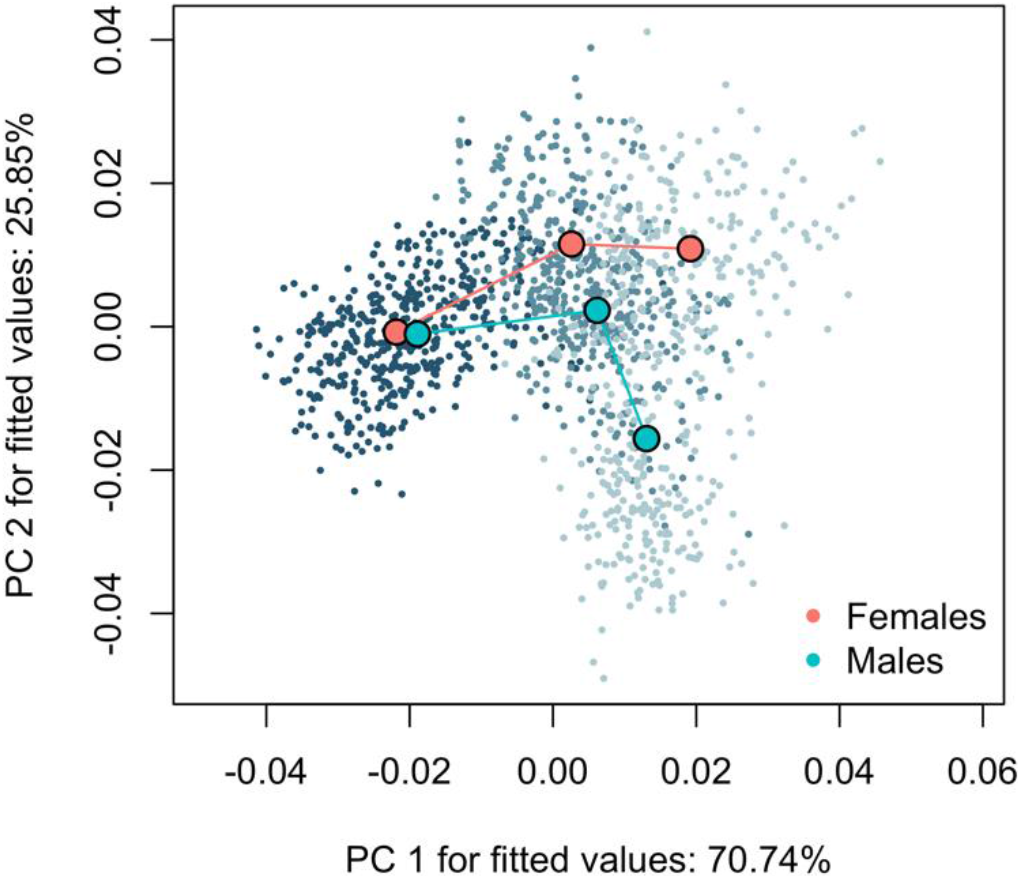
Phenotypic Trajectory Analysis onto the first two principal components based on the covariance matrix of group means. Each point represents one individual. Descending color intensity depicts the different time points (i.e. 18, 24 and 36). Large points represent mean shape estimates for each cross type at each trajectory point, and are connected in chronological order.

We focused on sexually mature individuals at month 36 to study the maximum dissimilarity in shape between males and females. The first principal component (**Fig. 5**) exhibited a clear distinction between sexes, although their location in the morphospace also overlapped. Despite cross having a significant effect on shape, its visualisation through examination of the first PCs did not show any apparent clustering (**Fig. 7D**). Thus shape changes along PC1 depicted male-female shape differentiation, which occurred mostly on the relative depth and length of the head: males showed substantially larger heads compared to females, deeper bodies (on the hump and pelvic fin area) and slightly shorter caudal peduncles. Additionally, males seem to present pointier snouts compared to females, but no differences in the position of upper and lower jaw were detected. We also examined whether pure crosses or reciprocal hybrids clustered towards the “most male or female” morphospace, but found no clear structure (SM5, Fig. S5 for visualisation).

**Figure 5.**
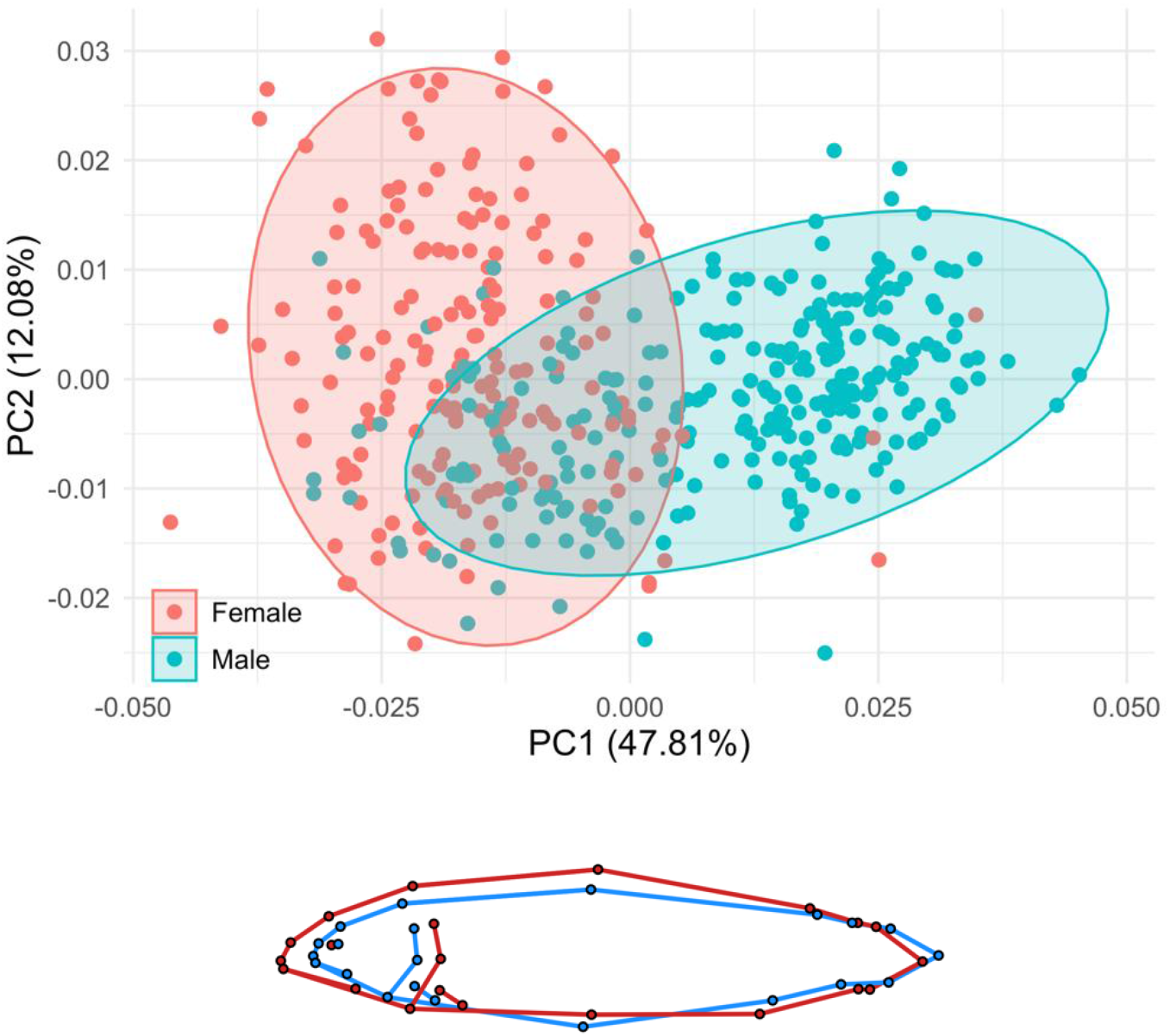
Principal component analysis of Procrustes coordinates of sexually mature specimens at month 36. Each point represents one individual. Females are depicted in salmon color, whereas males in blue. Shaded, 95% confidence ellipses for each sex. The wireframe represents shape changes across PCs. Blue and red wireframe depicts minimum and maximum values of each principal component.

### Morphs and hybrids differ in their ontogenetic trajectories

As previously stated, there was a significant effect of *cross* in both data sets and significant pairwise differences in means between all cross types. Taking into account that we pooled together different ages, sexes and sexual maturation states of fish with a very recent evolutionary history grown in common garden conditions, having a signal for *cross* indicates that traits traditionally associated with benthic-limnetic adaptations on Arctic charr have a strong genetic component. We then asked whether ontogenetic trajectories between the two morphs differed, what the position of the hybrids was and whether there were signatures for canalisation of traits associated to benthic-limnetic adaptation.

PTA showed overall differences in ontogenetic trajectories between each cross type (**Fig. 6**). We found highly significant differences in trajectory location between all pairs (p = 0.001), direction (Pr angle ≤ 0.007 for each pair), and shape (Pr > d = 0.001 for each pair). However, only significant differences in amount of shape change (i.e., absolute path trajectory lengths) between both pure crosses and PLxSB (Pr > d = 0.007 and 0.004 respectively) were found, showing PLxSB having the longest trajectory (maximum absolute path distance = 0.0672).

**Figure 6.**
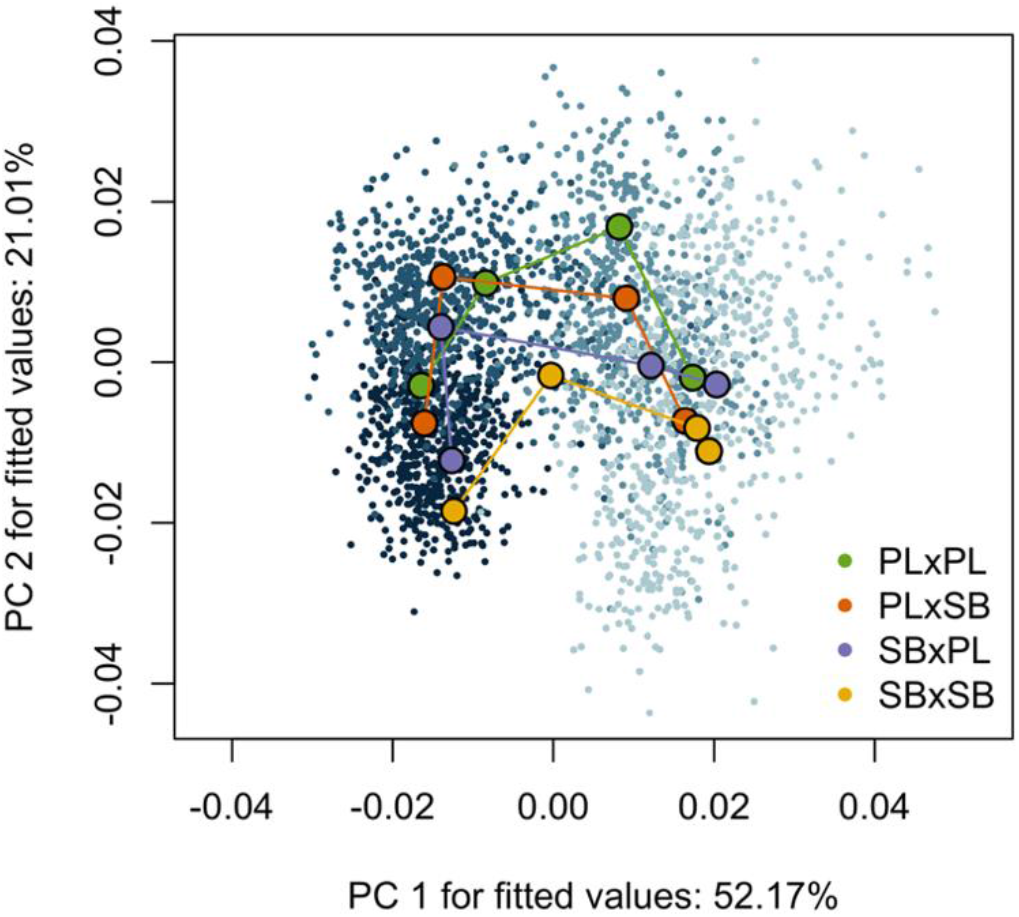
Phenotypic Trajectory Analysis onto the first two principal components based on the covariance matrix of group means. Each point represents one individual. Descending color intensity depicts the different time points as in Figure 2. Large points estimate mean shapes for each cross type at each trajectory point and are connected in chronological order.

When plotted over the PCA, the different cross types started their mean trajectories at similar values of PC1, slightly differing along PC2, where pure crosses PL and SB showed divergent phenotypes and reciprocal hybrids occupied an intermediate position. Interestingly, the trajectories of pure crosses developed parallelly towards month 18, whereas hybrids moved to higher values of PC2 and closer to pure PL. Trajectories of pure crosses stopped being parallel from month 18 onwards and showed their largest morphological difference at month 24. On the other hand, hybrids advanced in a similar fashion towards month 24, where they reached similar mean values of PC1 than their parental species but occupied an intermediate space in PC2. At month 36, cross type means converged in both PC1 and PC2, although morphological disparity at this time point increased, likely driven by -at least- individuals who reached sexual maturation and showed secondary sexual traits (SM4, Table S4, Fig. S4.2).

### The effect of cross type varies during ontogeny

For each of the four life stages, MANOVAs showed significant effects of *log(Csize), cross* and the interactions *cross:family* and *log(Csize):cross:family* (except the later was not significant at month 12) (**Table 3**). At month 12, the effect of *cross* alone was present and had a strong signal. Although its effect size (Z) remained relatively constant through time, the amount of variance explained by *cross* alone increased from 11% at 12 months towards 19.7% a year later. Interestingly, and as expected in the view of previous results, its R2 decreased substantially at month 36 (4.4%) and its Z score value was only a half relative to previous months. The effect of nested families was highly significant at each time point, indicating that shape variation among families is meaningful compared to cross type effect. Further, the relevance of *log(Csize)* and *log(Csize):cross:family* was not dismissible, even though fish were phenotyped on the same day for each sampling month (except for month 18, when pure crosses were phenotyped only 15 days after the hybrids).

**Table 3.** Covariates effects from independent MANOVAs at for months 12, 18, 24 and 36, using residual randomization. Error terms updated to account for nested families within cross types.

|  | Month 12 |  |  |  | Month 18 |  |  |  |
| --- | --- | --- | --- | --- | --- | --- | --- | --- |
|  | Rsq | Z | pr (>F) |  | Rsq | Z | pr (>F) |  |
| $\log(Csize)$ | 0.037 | 8.136 | 0.001 | ** | 0.080 | 10.191 | 0.001 | ** |
| $cross$ | 0.106 | 4.374 | 0.001 | ** | 0.131 | 5.310 | 0.001 | ** |
| $cross:family$ | 0.110 | 13.774 | 0.001 | ** | 0.097 | 14.776 | 0.001 | ** |
| $\log(Csize):cross$ | 0.020 | -0.831 | 0.788 | | 0.023 | 0.360 | 0.037 | |
| $\log(Csize):cross:family$ | 0.016 | 0.596 | 0.283 | | 0.016 | 1.872 | 0.036 | * |
| Residuals | 0.675 |  |  |  | 0.542 |  |  |  |

|  | Month 24 |  |  |  | Month 36 |  |  |  |
| --- | --- | --- | --- | --- | --- | --- | --- | --- |
|  | Rsq | Z | pr (>F) |  | Rsq | Z | pr (>F) |  |
| $\log(Csize)$ | 0.042 | 8.229 | 0.001 | ** | 0.082 | 7.091 | 0.001 | ** |
| $cross$ | 0.197 | 4.737 | 0.001 | ** | 0.044 | 2.618 | 0.008 | ** |
| $cross:family$ | 0.006 | 14.375 | 0.001 | ** | 0.071 | 7.495 | 0.001 | ** |
| $\log(Csize):cross$ | 0.139 | -4.823 | 1.000 | | 0.009 | -1.241 | 0.899 | |
| $\log(Csize):cross:family$ | 0.023 | 3.331 | 0.002 | ** | 0.037 | 4.172 | 0.001 | ** |
| Residuals | 0.574 |  |  |  | 0.727 |  |  |  |

### The directionality of the cross partially explains differences between hybrids

The first two components of the PCA explained a great proportion of the variation and showed group clustering in different fashions at each different time point (**Fig. 7**), spanning the range of possibilities that theory predicts about hybrid phenotypes (i.e., intermediate, parent-like or transgressive). Regarding allometry, only in month 12 the null hypothesis of homogeneity of slopes between cross types was not rejected (p = 1.000). However, we decided not to correct for common allometry to simplify result interpretation by making time points comparable. Furthermore, specimens were phenotyped at the time points within one day (except for month 18, when pure crosses were photographed only 15 days later than the hybrids), and although there were differences in size between cross types at certain time points, overall size variation was very small (SM1, Table S2.1).

**Figure 7.**
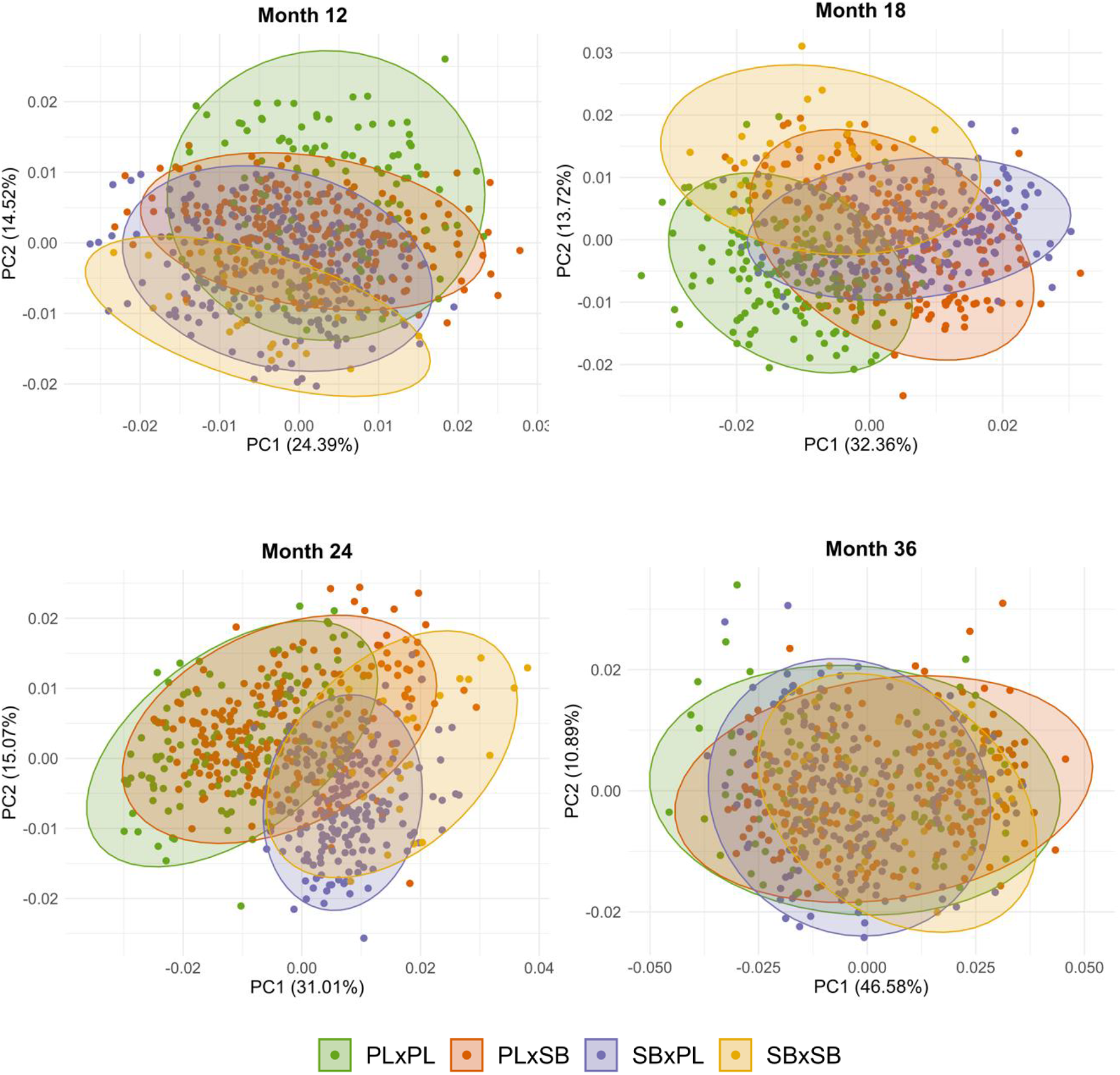
Two first principal components of separate principal component analyses of months 12, 18, 24 and 36. Each point represents one individual and shaded areas depict 95% confidence ellipses.

We found significant differences between all pairs of cross types within each time point (p<0.05) when fitting explicit reduced models (*shape ~ 1 + log(Csize)*) (see pairwise comparisons in SM6). At month 12 (**Fig. 7A**), cross types appeared to be considerably scattered in the morphospace. However, 95% confidence ellipses around each cross means allowed the visualisation of pure crosses occupying more extreme positions along PC2 (16.14%), whereas reciprocal hybrids laid in the middle, but they did not completely overlap with each other. At month 18 (**Fig. 7B**), both PC1 (29.95%) and PC2 (13.34%) explained shape differences between morphs. This time, a substantial proportion of hybrids showed transgressive (i.e., outside of the parental range) phenotypes at high values of PC1, the main axis of variance (29.95%). The rest of the hybrid specimens laid between their parental phenotypes. This pattern switched again at month 24 (**Fig. 7C**), where hybrid shape variation was mostly contained within their maternal morphospace. Nevertheless, the positioning of reciprocal hybrids was not symmetrical: SBxPL shape variability is particularly reduced compared to PLxSB, which completely overlapped with PLxPL, and also expanded towards SBxSB and SBxPL’s morphospaces. No clear cross type clustering was found at month 36 (**Fig. 7D**) when examining the first ten principal components, despite overall shape differences were significant between all pairs (SM6, table S4). Wireframes of predicted landmark configurations (SM7) at minimum and maximum PC values showed mostly changes in body depth, relative head size, caudal peduncle length, position of the pectoral fin and relative length of the upper and lower jaw.

## DISCUSSION

We found extensive differences in shape variation across time in common garden conditions, showing a strong genetic component driving shape differences between time points, sexes and contrasting ecomorphs. Our results indicated that growth is the main driver of shape variation across time and provided evidence for a strongly genetically-controlled ontogenetic shift that gives rise to the limnetic morph. Before month 36, shape variation differed between morphs, and reciprocal hybrids tended to group towards their respective maternal phenotypes. Additionally, the onset of sexual maturation triggers differences in sex ontogenetic trajectories and shape variation at different time points, likely dissipating the canalisation of traits traditionally associated with benthic-limnetic adaptation. The interplay between traits associated with benthic-limnetic adaptation and sexual dimorphism is complex and dynamic during ontogeny.

### The ontogeny of benthic-limnetic traits

Growth was one of the main drivers of shape across time in common garden conditions, justifying subsequent examination of ontogenetic trajectories in different groups. As hypothesized, individuals at extreme time points (12 and 36 months old) showed the most distinct shapes. Younger specimens presented phenotypes commonly associated with bottom-feeding morphologies: rounded snouts and subterminal jaws, to optimise food suction on the benthos, and a large head relative to their body. Specimens at month 36 showed more pelagic-like forms: pointed snouts with terminal jaws, enabling them to catch floating particles, and relatively smaller heads. This result is consistent with the previous literature, since Arctic charr ecomorphs -also across lakes-start feeding on the benthos at juvenile stages. Benthic morphs retain these juvenile-like traits as adults, whilst limnetic morphs migrate towards pelagic areas in order to feed on plankton (planktivorous morph, PL) or other fish (piscivorous, PI) (Skúlason et al., 1989). During the experimental setup, food was administrated on the surface, sank down and stayed on the bottom of the bucket, excluding the possibility that the benthic-limnetic feeding shift was environmentally induced. Yet, shape differences across time were still distinguishable, and provided evidence that this ontogenetic shift is genetically controlled, although it may be exacerbated when fish are exposed to distinct environmental conditions.

The retention of juvenile traits in adult stages is a type of heterochrony called paedomorphosis. Previous studies in common garden of different ecomorphs from Thingvallavatn (Skúlason et al., 1989, Parsons et al., 2011, Kristjánsson et al., 2018) either showed evidence or speculated that heterochronic changes are the source of phenotypic variation underlying differences between ecomorphs. To our knowledge, this is to the only common-garden experiment showing paedomorphosis on Arctic charr in advanced ontogenetic time points. This means that the ontogenetic repatterning of traits associated with benthic-limnetic adaptations spans a considerable time period of charr ontogeny and should be taken into account when studying these traits in mature individuals. We therefore would have theoretically expected a shorter trajectory length (as a proxy of a “lower amount” of shape change) in SB compared to PL. However, the low sample size of SB, the heterogeneous onset of sexual maturation within the sampled individuals and other developmental processes might have influenced this result. In any case, environmental conditions likely emphasize the process of ontogenetic repatterning as a source for phenotypic variation to generate contrasting morphs (Parsons et al., 2011, Kristjánsson et al., 2018).

Additionally, we asked whether traits associated with benthic-limnetic adaptations are present during ontogeny in the different ecomorphs. Although the effect of cross type did not explain a substantial proportion of the variance in the main model, it was still latent despite shape variation attributed to ontogenetic growth and sexual dimorphism. This indicates that there is a genetic component underlying the development of multiple traits in contrasting ecomorphs. In fact, a recent study showed morphological differences between PL and SB morphs from Thingvallavatn even before first feeding in a common garden setup (Ponsioen 2020).

Moreover, support for genetic canalisation of putative adaptive traits was found in the increase of the relative load of cross type across time and the divergence of pure morph’s ontogenetic trajectories towards month 24. To confirm this, we expected morphological disparity within-groups to decrease, and between-groups to increase across time. Nevertheless, both within- and between- morphological disparity during the first three time points remained relatively constant, but substantially increased at month 36. This may have been due to differences associated with the onset of sexual dimorphism and the heterogeneity in its starting point within this dataset. The strong family effect and confounding environmental cues (since feeding was mainly benthic-like) may have additionally influenced this result.

Already Skúlason et al. (1989), in a common garden experiment on Thingvallavatn ecomorphs and their hybrids, showed an increase in frequency of extreme values of canonical scores during ontogeny in pure morphs, which supports trait canalisation. Additionally, LB and PL Arctic charr from Thingvallavatn grown in captivity under benthic and limnetic treatments showed increased trait canalisation from 90 to 160 days after initiation of diet treatments relative to less divergent morphs from another Icelandic smaller lake (Parsons et al., 2011). Although the latter was not a common garden experiment, the comparison between benthic and limnetic treatments on both lakes suggests that benthic-limnetic trait canalisation in Thingvallavatn is strongly genetically induced.

#### Hybrids are not that different after all

Hybrid‘s trajectories were different to their parental ecomorphs and between each other, meaning that developmental programs in hybrids function in a different fashion, although data imbalance due to low sample size in pure crosses, especially in SB individuals should be considered. However, at least in a common garden setup, there is no reason for different trajectories being attributed to their ability to feed and grow, as growth differences were similar among groups.

Before month 36, pure crosses tend to diverge across time, whereas hybrids show different patterns relative to their parental phenotypes. Most hybrids adopted their mother’s phenotype at month 24, and at month 36, major differences in shape variation did not occur between different cross types, but between sexes. Hence, there is no evidence pointing towards a decrease of hybrid’s viability, at least in terms of genetically driven shape differences and ontogenetic trajectories. This is consistent with recent literature on SB and PL morphs from Thingvallavatn at earlier life stages, where it has been shown that the trait covariance patterns in F1 hybrids tend to differ from pure crosses, rather than differences in several mean trait values (Horta-Lacueva et al., 2020, *in press*).

### The onset of sexual maturation

Our results indicate that the onset of sexual maturation plays a fundamental role in explaining shape variation. The effect of sex is already significant at month 18, but its importance seemingly increases during ontogeny until month 36. Accordingly, extensive differences in shape between males and females were found at this time point, indicating a strong genetic component of sexual dimorphism.

Sexual dimorphism within the *Salmonidae* family is clearly marked due to sexual selection (De Gaudemar, 1998). Classic examples of overly developed sexual traits in salmonids occur mostly in males, characterised by larger body size, pointier or even hooked snouts, exaggerated humps and bright colorations, as a result from high male density and competition (Fleming & Gross, 1996). These traits are in line with the shape changes observed at month 36 along the first principal component, in which males showed a pointier snout, bigger head and humped dorsal area. However, we did not find significant differences in centroid size between males and females and hence it can be argued that size differences may be environmentally induced, and it is independent of -at least-the sexual secondary traits which we are able to detect in this common garden setup. This is supported by the non-significant interaction between size and sex at months 18 and 36. In addition, the increase of morphological disparity of males relative to females may indicate that the development of secondary sexual traits in males is more heterogenous and not necessarily genetically canalised, and the mechanisms leading to it are complex (see e.g., Woram et al., 2003, Sutherland et al., 2019) and remain unclear.

#### Mitigation of benthic-limnetic adaptive traits

When looking at the combined effects of sexual dimorphism and traits associated to benthic-limnetic adaptations, we showed that their dynamic pattern is reversed. This means that, despite the effect of cross type and sex being significant at each separate time point, the relative load of cross decreases while sex increases towards month 36. These results are reinforced by both ecomorph and sex ontogenetic trajectories: mean shapes of different morphs converge at month 36, as morphological disparity dramatically increases, and sex ontogenetic trajectories diverge.

One explanation is that sexual maturation, developmental programs and the processes driving benthic-limnetic morphological divergence are decoupled, at least at certain time points and in the same environment. This is supported by the non-significant interaction between sex and cross in the main model during our experimental setup, indicating that the cross type and sex independently affect shape. Additionally, as it has been reported in other adaptive radiations, differences in certain adaptive traits can be incredibly subtle (e.g.: pumpkin sunfish (Jastrebski & Robinson, 2004); or female preference in Malawi cichlids (Ding et al., 2014)). A combination of both hypotheses may have facilitated the mitigation of putative benthic-limnetic adaptations during the onset of sexual maturation.

### The interplay between sexual dimorphism and adaptive divergence

We were not able to determine whether sexual dimorphism in this system is originated prior-, post-ecological diversification, or a combination of both, and studying sexual dimorphism within cross type was not possible due to data imbalance (especially in the case of SBxSB). Considering Arctic charr colonised different water bodies in Iceland and that it (1) parallelly evolved distinct ecomorphs, and that, (2) in principle, there are no cases of populations evolving ecological sexual dimorphism before ecological diversification, one can argue that what we observe is the result of ancestral sexual dimorphism and subsequent adaptive radiation. This is additionally supported by a widespread strong sexual dimorphism within the salmonid clade, hence sexual traits may be well conserved.

If sexual dimorphism arose after adaptive divergence, aspects of shape that differ between sexes should not be the same as those differing between ecomorphs, in order to maintain assortative mating (and unless assortative mating somehow occurred prior to phenotypic ecological diversification). In our study, major differences between sexes mainly correspond with head size relative to the body, body depth and snout shape. Nevertheless, the relationship between upper and lower jaw (subterminal jaws are associated with benthic feeding strategies, whereas terminal jaws are common when feeding elsewhere on the water column) seems to be proportional between males and females. Thus, differences between sexes only slightly overlap with differences observed between ecomorphs and may potentiate ecological adaptation within morphs in an additive fashion, as it has been shown to be the case in sticklebacks (Cooper et al., 2011). Looking at the sex-coloured PCA at month 36, we are not able to determine in which degree sex traits are confounded due to underlying adaptive-associated traits which do not lay at PCs extremes and thus their variation is not represented in Principal Component Analysis. Another confounding factor may be heterogeneous sexual maturation time (i.e., some individuals were not mature yet, and thus not sexed and not used for sexual dimorphism analysis) and/or other differences in life history traits between morphs (Jonsson 1988).

## CONCLUSION

We found that phenotypic traits associated with benthic-limnetic adaptations are present and are genetically controlled, not only throughout ontogeny, but also at different time points between PL, SB and their hybrids. Sexual maturation is key in this scenario, since developmental programs driving the onset of the breeding season may dissipate the canalisation for adaptive traits in similar environments at certain life stages. Nevertheless, the interplay during development between traits associated with ecological diversification and sexual maturation is complex and more efforts should be directed towards studying their relationship in this and other adaptive radiation systems.

## Acknowledgements

We thank Kári H. Árnason, Rakel Þorbjörnsdóttir, and Christian Beuvard for the maintenance of the experimental setup at the rearing facility at Verið, Sauðárkrókur (Hólar University College, Iceland). We also thank Prof Sigurður S. Snorrason for his helpful advice and comments.

## Authorship contribution

Marina de la Cámara conducted the analysis and wrote the manuscript. Lieke Ponsioen collected the data, phenotyped the specimens and critically revised the manuscript. Quentin J.B. Horta-Lacueva critically revised the manuscript. Kalina H. Kapralova conceived the study, established the crossing design, reared the embryos, collected the data and contributed to the writing of the manuscript. All authors gave their final approval for publication and agree to be accountable for the work therein.

## Ethical note

The rearing and the experimental work was conducted in the facilities of Hólar University Aquaculture Research Station, which has an operational license under the Icelandic Aquaculture law (Law No. 71/2018). This law includes clauses of best practices for animal care and experimental work. Decisions on the sample size and on the design of the common-garden experiment were made to ensure that additional studies could be conducted with data collected on the same specimens.

## Data accessibility

The data will be deposited onto the Dryad Digital Repository upon acceptance.

