## Supplemental file for "The dynamic ontogenetic patterns of adaptive divergence and sexual dimorphism in Arctic charr"

#### SUPPLEMENTARY MATERIAL

### SM1. Crossing design

**Table S1.** Crossing desgin with number of individuals and included in final analyses per family and time point.

| cross | familiy | # ind | | | |
|---|---|---|---|---|---|
| | | month 12 | month 18 | month 24 | month 36 |
| **PLxPL** | H1509 | 35 | 30 | 27 | 27 |
| | H1513 | 35 | 30 | 26 | 25 |
| | H1516 | 30 | 29 | 29 | 26 |
| | H1517 | 30 | - | - | - |
| | H1518 | 14 | 9 | 10 | 9 |
| | H1525 | - | 52 | 37 | 35 |
| | **6** | 144 | 150 | 129 | 122 |
| **PLxSB** | H1512 | 28 | 14 | 13 | 14 |
| | H1514 | 23 | 20 | 22 | 22 |
| | H1520 | 3 | 2 | 3 | 3 |
| | H1524 | 29 | 29 | 27 | 28 |
| | H1526 | 92 | 93 | 84 | 87 |
| | H1527 | 54 | 48 | 41 | 43 |
| | H1531 | 32 | 22 | 25 | 25 |
| | **7** | 261 | 228 | 215 | 222 |
| **SBxPL** | H1505 | 24 | 17 | 16 | 15 |
| | H1506 | 101 | 107 | 94 | 96 |
| | H1507 | 72 | 76 | 76 | 72 |
| | H1508 | 33 | 33 | 31 | 33 |
| | **4** | 230 | 233 | 217 | 216 |
| **SBXSB** | H1502 | 4 | 16 | 16 | 15 |
| | H1503 | 22 | 17 | 17 | 15 |
| | **2** | 26 | 33 | 33 | 30 |
| | **19** | | | | |

### SM2. Centroid size

**Table S2.1.** Centroid size summary by *month:cross*.

| month | cross | Csize mean (cm) | std. dev (cm) |
|---|---|---|---|
| **12** | PLxPL | 8.94 | ± 0.94 |
| | PLxSB | 8.86 | ± 0.67 |
| | SBxPL | 9.33 | ± 0.84 |
| | SBxSB | 9.35 | ± 0.84 |
| **18** | PLxPL | 13.15 | ± 1.49 |
| | PLxSB | 11.82 | ± 1.47 |
| | SBxPL | 11.88 | ± 1.47 |
| | SBxSB | 13.71 | ± 1.84 |
| **24** | PLxPL | 18.81 | ± 2.45 |
| | PLxSB | 18.52 | ± 2.16 |
| | SBxPL | 18.58 | ± 2.13 |
| | SBxSB | 19.62 | ± 2.72 |
| **36** | PLxPL | 25.46 | ± 4.75 |
| | PLxSB | 24.90 | ± 4.53 |
| | SBxPL | 25.57 | ± 3.31 |
| | SBxSB | 23.35 | ± 3.50 |

**Table S2.2**. Centroid size summary by *month:sex*.

| month | sex | Csize mean (cm) | std. dev (cm) |
|---|---|---|---|
| **12** | NA | 9.07 | ± 0.82 |
| **18** | f | 12.08 | ± 1.58 |
| | m | 12.57 | ± 1.68 |
| | NA | 11.95 | ± 1.51 |
| **24** | f | 18.70 | ± 2.19 |
| | m | 19.19 | ± 2.05 |
| | NA | 17.69 | ± 2.39 |
| **36** | f | 25.87 | ± 3.58 |
| | m | 25.41 | ± 4.29 |
| | NA | 23.83 | ± 4.30 |

#### SM3. MANCOVA with the full dataset

**Table S3**. Table 2. Final mixed-model MANOVA using residual randomization on shape of the full dataset of individuals phenotyped at 12, 18, 24 and 36 months after hatching. Error terms updated to account for nested families within cross types.

| | d.f. | SS | MS | Rsq | F | Z | Pr (>F) | |
|---|---|---|---|---|---|---|---|---|
| *log(Csize)* | 1 | 0.528 | 0.5275 | 0.262 | 1403.14 | 12.60 | 0.0010 | ** |
| *month* | 3 | 0.257 | 0.0855 | 0.128 | 227.52 | 16.02 | 0.0010 | ** |
| *cross* | 3 | 0.090 | 0.0299 | 0.045 | 4.22 | 4.30 | 0.0010 | ** |
| *log(Csize):month* | 3 | 0.022 | 0.0074 | 0.011 | 19.71 | 10.42 | 0.0010 | ** |
| *log(Csize):cross* | 3 | 0.017 | 0.0056 | 0.008 | 0.78 | -0.71 | 0.7560 | |
| *cross:family* | 14 | 0.099 | 0.0071 | 0.049 | 18.83 | 18.02 | 0.0010 | ** |
| *log(Csize):month: cross:family* | 64 | 0.109 | 0.0017 | 0.054 | 4.53 | 18.84 | 0.0010 | ** |
| Residuals | 2367 | 0.890 | 0.0004 | 0.443 | | | | |
| Total | 2458 | 2.011 | | | | | | |

#### SM4. Morphological disparity

**Table S4.** Morphological disparity from Phenotypic Trajectory Analysis per sex and cross type at different time points.

| time point (month) | morphological disparity within time point | sex | morphological disparity within month:sex | cross type | morphological disparity within month:cross |
|---|---|---|---|---|---|
| 12 | $2.747 \times 10^{-4}$ | f | - | PLxPL | $2.507 \times 10^{-4}$ |
| | | | | PLxSB | $2.781 \times 10^{-4}$ |
| | | m | - | SBxPL | $2.730 \times 10^{-4}$ |
| | | | | SBxSB | $2.382 \times 10^{-4}$ |
| 18 | $2.722 \times 10^{-4}$ | f | $2.965 \times 10^{-4}$ | PLxPL | $3.240 \times 10^{-4}$ |
| | | | | PLxSB | $3.091 \times 10^{-4}$ |
| | | m | $2.995 \times 10^{-4}$ | SBxPL | $3.212 \times 10^{-4}$ |
| | | | | SBxSB | $3.002 \times 10^{-4}$ |
| 24 | $3.179 \times 10^{-4}$ | f | $2.878 \times 10^{-4}$ | PLxPL | $3.700 \times 10^{-4}$ |
| | | | | PLxSB | $3.066 \times 10^{-4}$ |
| | | m | $3.993 \times 10^{-4}$ | SBxPL | $2.414 \times 10^{-4}$ |
| | | | | SBxSB | $3.222 \times 10^{-4}$ |
| 36 | $5.974 \times 10^{-4}$ | f | $4.342 \times 10^{-4}$ | PLxPL | $5.830 \times 10^{-4}$ |
| | | | | PLxSB | $6.136 \times 10^{-4}$ |
| | | m | $5.374 \times 10^{-4}$ | SBxPL | $5.377 \times 10^{-4}$ |
| | | | | SBxSB | $5.488 \times 10^{-4}$ |

**Fig. S4.1.** Morphological disparity across time in males (blue) and females (red).

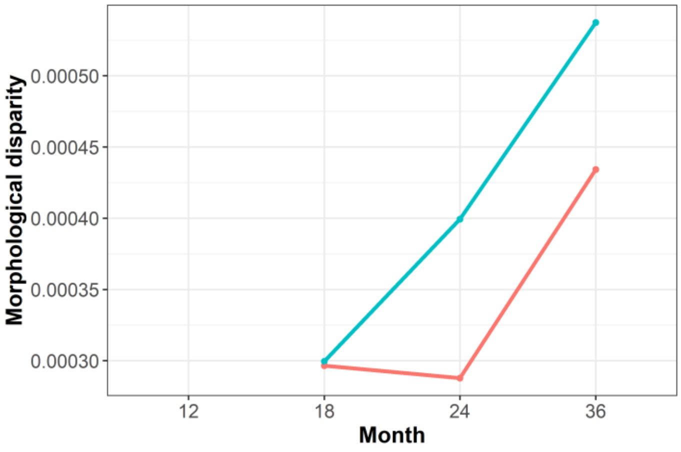


**Fig S4.2.** Morphological disparity across time in the different morphs

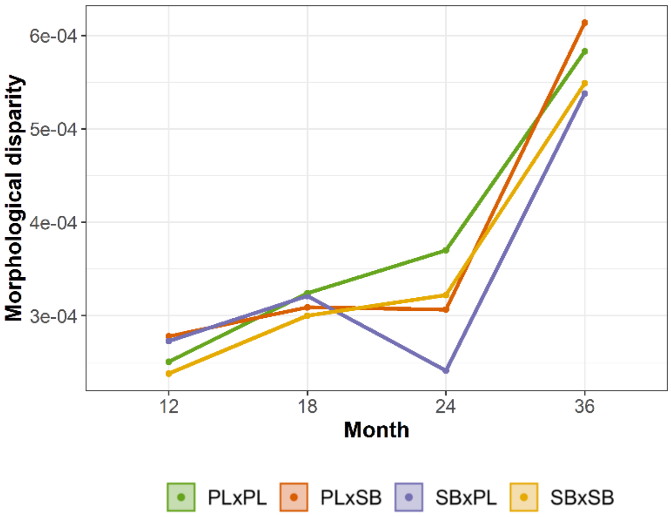

#### SM5. PCAs at month 36

**Figure S5.** Two first principal components of month 36, showing that either pure crosses or reciprocal hybrids did not cluster towards the "most male or female" area of the morphospace. Each point represents one individual and shaded areas depict 95% confidence ellipses.

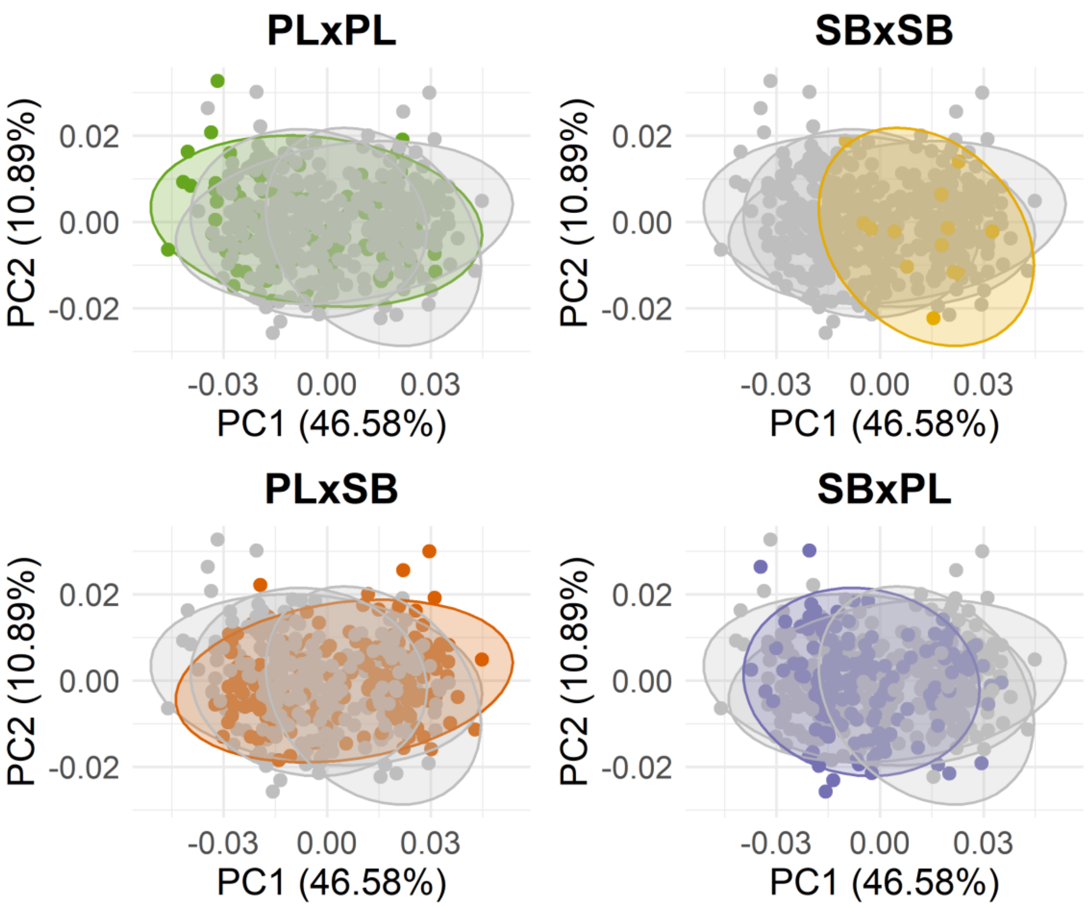

**SM 6.** Pairwise comparisons in means and variances between cross types at each time point, using 1000 permutations

#### **Table S6.1.** Month 12

| **Pairwise distances between means, plus statistics** | | | | |
|---|---|---|---|---|
| | d | UCL(95%) | Z | Pr>d |
| PLxPL:PLxSB | 0.0093 | 0.0047 | 7.493 | 0.001 |
| PLxPL:SBxPL | 0.0145 | 0.0048 | 13.191 | 0.001 |
| PLxPL:SBxSB | 0.0214 | 0.0115 | 7.017 | 0.001 |
| PLxSB:SBxPL | 0.0096 | 0.0033 | 12.815 | 0.001 |
| PLxSB:SBxSB | 0.0174 | 0.0109 | 5.456 | 0.001 |
| SBxPL:SBxSB | 0.0101 | 0.0111 | 1.191 | 0.113 |

| **Observed variances by group** | |
|---|---|
| PLxPL | 0.000251 |
| PLxSB | 0.000278 |
| SBxPL | 0.000273 |
| SBxSB | 0.000238 |

| **Pairwise distances between variances, plus statistics** | | | | |
|---|---|---|---|---|
| | d | UCL(95%) | Z | Pr>d |
| PLxPL:PLxSB | 2.74E-05 | 3.055E-05 | 1.581 | 0.088 |
| PLxPL:SBxPL | 2.23E-05 | 3.103E-05 | 1.056 | 0.165 |
| PLxPL:SBxSB | 1.25E-05 | 5.938E-05 | -0.636 | 0.691 |
| PLxSB:SBxPL | 5.12E-06 | 2.476E-05 | -0.639 | 0.673 |
| PLxSB:SBxSB | 3.99E-05 | 5.744E-05 | 0.973 | 0.159 |
| SBxPL:SBxSB | 3.48E-05 | 5.664E-05 | 0.650 | 0.23 |

#### **Table S6.2.** Month 18

| **Pairwise distances between means, plus statistics** | | | | |
|---|---|---|---|---|
| | d | UCL(95%) | Z | Pr>d |
| PLxPL:PLxSB | 0.0140 | 0.0039 | 15.4466 | 0.001 |
| PLxPL:SBxPL | 0.0179 | 0.0037 | 20.2788 | 0.001 |
| PLxPL:SBxSB | 0.0253 | 0.0077 | 13.7834 | 0.001 |
| PLxSB:SBxPL | 0.0101 | 0.0037 | 11.6984 | 0.001 |
| PLxSB:SBxSB | 0.0203 | 0.0079 | 10.6856 | 0.001 |
| SBxPL:SBxSB | 0.0174 | 0.0078 | 9.0448 | 0.001 |

| **Observed variances by group** | |
|---|---|
| PLxPL | 0.000251 |
| PLxSB | 0.000257 |
| SBxPL | 0.000279 |
| SBxSB | 0.000236 |

| **Pairwise distances between variances, plus statistics** | | | | |
|---|---|---|---|---|
| | d | UCL(95%) | Z | Pr>d |
| PLxPL:PLxSB | 6.37E-06 | 2.87E-05 | -0.556 | 0.644 |
| PLxPL:SBxPL | 2.82E-05 | 2.78E-05 | 1.929 | 0.043 |
| PLxPL:SBxSB | 1.49E-05 | 5.11E-05 | -0.392 | 0.575 |
| PLxSB:SBxPL | 2.19E-05 | 2.41E-05 | 1.548 | 0.082 |
| PLxSB:SBxSB | 2.12E-05 | 4.96E-05 | 0.079 | 0.399 |
| SBxPL:SBxSB | 4.31E-05 | 4.99E-05 | 1.515 | 0.089 |

**Table S6.3.** Month 24

| Pairwise distances between means, plus statistics | | | | |
|---|---|---|---|---|
| | d | UCL(95%) | Z | Pr>d |
| PLxPL:PLxSB | 0.0120 | 0.0042 | 11.4171 | 0.001 |
| PLxPL:SBxPL | 0.0212 | 0.0039 | 20.3373 | 0.001 |
| PLxPL:SBxSB | 0.0302 | 0.0076 | 15.8586 | 0.001 |
| PLxSB:SBxPL | 0.0153 | 0.0037 | 17.1499 | 0.001 |
| PLxSB:SBxSB | 0.0219 | 0.0074 | 12.1959 | 0.001 |
| SBxPL:SBxSB | 0.0151 | 0.0072 | 7.6740 | 0.001 |

| Observed variances by group | |
|---|---|
| PLxPL | 0.000370 |
| PLxSB | 0.000307 |
| SBxPL | 0.000241 |
| SBxSB | 0.000322 |

| Pairwise distances between variances, plus statistics | | | | |
|---|---|---|---|---|
| | d | UCL(95%) | Z | Pr>d |
| PLxPL:PLxSB | 6.35E-05 | 3.78E-05 | 4.121 | 0.001 |
| PLxPL:SBxPL | 1.29E-04 | 3.92E-05 | 8.890 | 0.001 |
| PLxPL:SBxSB | 4.78E-05 | 7.20E-05 | 0.923 | 0.166 |
| PLxSB:SBxPL | 6.52E-05 | 3.54E-05 | 4.723 | 0.001 |
| PLxSB:SBxSB | 1.57E-05 | 6.92E-05 | -0.553 | 0.641 |
| SBxPL:SBxSB | 8.09E-05 | 6.82E-05 | 2.600 | 0.020 |

**Table S6.4.** Month 36.

| Pairwise distances between means, plus statistics | | | | |
|---|---|---|---|---|
| | d | UCL(95%) | Z | Pr>d |
| PLxPL:PLxSB | 0.0100 | 0.0056 | 6.0334 | 0.001 |
| PLxPL:SBxPL | 0.0104 | 0.0055 | 6.6022 | 0.001 |
| PLxPL:SBxSB | 0.0166 | 0.0109 | 4.5607 | 0.002 |
| PLxSB:SBxPL | 0.0105 | 0.0044 | 9.1559 | 0.001 |
| PLxSB:SBxSB | 0.0121 | 0.0105 | 2.7498 | 0.018 |
| SBxPL:SBxSB | 0.0124 | 0.0104 | 3.0227 | 0.010 |

| Observed variances by group | |
|---|---|
| PLxPL | 0.000583 |
| PLxSB | 0.000614 |
| SBxPL | 0.000538 |
| SBxSB | 0.000548 |

| Pairwise distances between variances, plus statistics | | | | |
|---|---|---|---|---|
| | d | UCL(95%) | Z | Pr>d |
| PLxPL:PLxSB | 3.15E-05 | 7.84E-05 | -0.013 | 0.443 |
| PLxPL:SBxPL | 4.47E-05 | 7.68E-05 | 0.625 | 0.244 |
| PLxPL:SBxSB | 3.44E-05 | 1.45E-04 | -0.531 | 0.64 |
| PLxSB:SBxPL | 7.62E-05 | 6.34E-05 | 2.552 | 0.017 |
| PLxSB:SBxSB | 6.59E-05 | 1.38E-04 | 0.233 | 0.346 |
| SBxPL:SBxSB | 1.03E-05 | 1.38E-04 | -1.054 | 0.873 |

**SM7.** Wireframes corresponding to Fig 7. Wireframes represent predicted shapes at each extreme of the two first principal components per time point. Low values of each principal component are represented by blue wireframes and high values by red wireframes.

Month 12 PC1

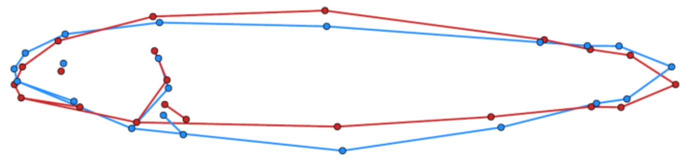

Month 12 PC2

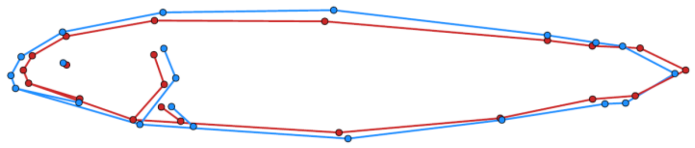

Month 18 PC1

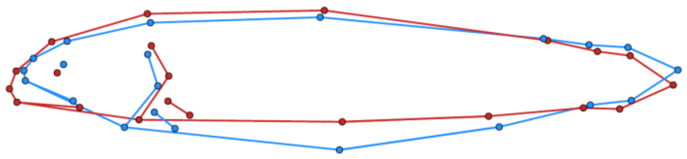

Month 18 PC2

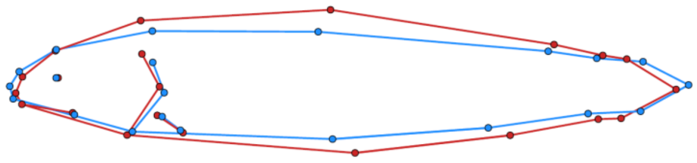

Month 24 PC1

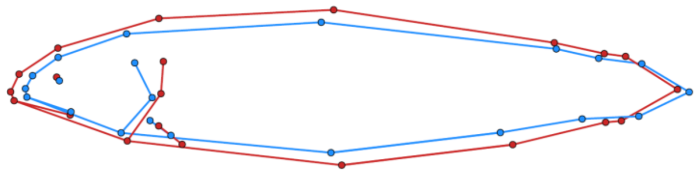

Month 24 PC2

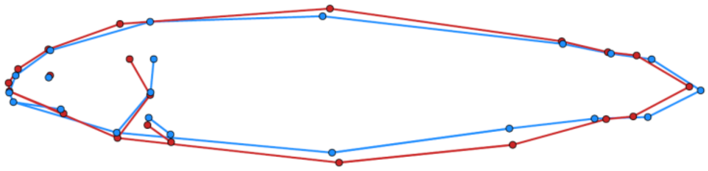

Month 36 PC1

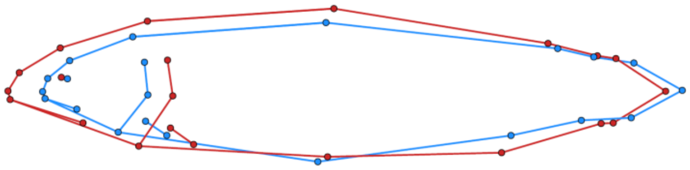
